# An Aurora Kinase-Dependent Role for Vif in Regulating HIV-1-Induced Cell-Cell Fusion

**DOI:** 10.64898/2026.06.15.732372

**Authors:** Samantha J. Tafrate, Brickley Littell, Jon P. Girard, Emily R. Mynar, Madeline K. Carr, Markus Thali, Menelaos Symeonides

## Abstract

Viral infectivity factor (Vif) is an HIV-1 accessory protein best known for its counteraction of APOBEC3 enzymes, interferon-inducible host defenses against viral infection, as well as PPP2R5A-E, which are regulatory subunits of the PP2A cellular phosphatase holoenzyme, resulting in striking Vif-dependent phosphoproteome remodeling. One reported consequence of this remodeling is hyperphosphorylation of several Aurora kinase substrates in HIV-1 infected cells, which is reversed when Vif-deficient virus is used. We previously showed that infection of T cells with Vif-deficient HIV-1 results in significantly accelerated formation of syncytia compared to wild-type HIV-1 infection. More recently, others have shown that application of Aurora kinase B inhibitors during HIV-1 infection in T cells also results in a similar hyperfusogenic phenotype. Both effects were specific to Env-driven cell-cell fusion, and did not influence virus infectivity. We thus hypothesized that Vif’s influence on the rate of HIV-1-induced cell-cell fusion was mediated by Aurora kinase activity.

To start testing this hypothesis, we have evaluated the effects of a small panel of Aurora kinase inhibitors on HIV-1-induced cell-cell fusion in the presence or absence of Vif. Our results replicate the previously documented increase in cell-cell fusion in the absence of Vif, as well as the increase in cell-cell fusion observed upon inhibition of Aurora kinase B in the presence of Vif. Critically, we now present evidence that Vif deletion significantly blunts the impact of Aurora kinase inhibition on cell-cell fusion, supporting our hypothesis that Vif-mediated regulation of cell-cell fusion depends on Aurora kinase signaling dysregulation, likely because of PPP2R5A-E degradation. Further, we document that the cell-cell fusion regulator downstream of Aurora kinase signaling is likely Ezrin, which we have previously shown to prevent excess HIV-1-induced syncytium formation when in its phosphorylated (activated) state. Taken together, these findings establish a Vif, Aurora kinase, and Ezrin-dependent mechanistic framework for the regulation of HIV-1-induced cell-cell fusion in infected T cells which likely helps preserve optimal cell-to-cell virus transmission.

## 1. Introduction

Vif (Viral infectivity factor) is an accessory protein encoded by human immunodeficiency virus type 1 (HIV-1). Vif is required for production of infectious virions in primary CD4^+^ T cells, macrophages, and commonly used cell lines ^1^, primarily because it protects the viral genome from the host restriction factor apolipoprotein B mRNA-editing catalytic polypeptide (APOBEC3). APOBEC3 is a family of DNA cytosine deaminases that cause C to U deamination of the viral cDNA during reverse transcription ^2^. Vif prevents APOBEC3s from inducing lethal mutations in the viral genome by mediating proteasomal degradation of several (but not all) APOBEC3s before they can be encapsidated in the virion ^3^. Vif induces heterodimerization of CBFβ, then recruitment of Elongin B and C, Cul5, and RBX2 to form the E3 ubiquitin ligase complex which polyubiquitinates ABOPEC3s, targeting the proteins for destruction^1^. The action of APOBEC3s is not perfectly counteracted by Vif; low level of expression sufficiently protects against lethal mutagenesis yet allows for a degree of APOBEC3-induced mutations that contribute to the genetic diversity of the viral genome.

In addition to its role in counteracting APOBEC3s, Vif has been known to be involved in G2/M cell cycle arrest since the early 2000s, but the exact mechanism of involvement was not understood until more recently. G2/M cell cycle arrest, i.e. stalling of progression from the G2 phase of the cell cycle to the M phase, is a highly regulated cellular process that can be halted in response to activation of the DNA damage checkpoint or DNA replication checkpoint ^4^. Early studies showed that Vif was sufficient to induce G2/M cell cycle arrest in multiple cell types, and that this function was independent of APOBEC degradation ^5^. The HIV-1 accessory protein Vpr also independently induces G2/M cell cycle arrest ^6–8^. It was recently shown that the G2/M arrest induced by Vif may be significantly more cytotoxic than the one induced by Vpr, and that Vpr thus acts to preserve G2/M arrest (which helps amplify virus production through provirus duplication) while “protecting” the infected cell from the toxic effects of Vif ^9^.

Vif mutants defective in binding Cul5 and Elongin C/B, as well as depletion of CBFβ, rendered Vif unable to induce cell cycle arrest ^1^. Proteomic analysis of HIV-1 infection with and without Vif revealed depletion of components of the protein phosphatase 2A holoenzyme (PP2A), specifically the five PP2A-B56 regulatory subunits, PPP2R5A-E^10^. PP2A is a ubiquitous serine/threonine protein phosphatase involved in multiple cellular processes including cell cycle regulation and the G2/M transition ^10,11^. Vif accomplishes this by forming a Vif–CBFβ–Cul5 E3 ligase–PPP2R5A complex, allowing for polyubiquitination of the regulatory subunit by Cul5 E3 ligase, as well as directly blocking the PP2R5A substrate binding pocket ^11^. PPP2R5A-E depletion is a conserved Vif function across phylogenetically distant lentiviruses, suggesting that this function could pose a selective advantage for HIV-1 ^10^. Vif-mediated degradation of PPP2R5A-E results in rapid global changes in the host cell phosphoproteome ^10,12,13^, including increased phosphorylation of proteins involved in cell-cycle regulation and activation of the DNA damage response ^10^. Vif mutants incapable of PPP2R5 degradation no longer induce cell cycle arrest, establishing a mechanism for Vif-mediated G2/M arrest ^1,10^. Altogether, these findings support the idea that Vif-mediated regulation of PP2A activity and host phosphoproteome remodeling are important for securing efficient HIV-1 replication.

While multiple cellular pathways are dysregulated as a result of Vif-mediated PPP2R5A-E depletion, one of the most significant Vif-driven changes in the phosphoproteome is the activation of Aurora kinases (AURKs) ^10^. Aurora kinases A, B, and C (AURKA/B/C) are a family of serine/threonine kinases involved in the regulation and progression of mitosis. The Aurora kinase family consists of three closely related proteins: Aurora kinase A (AURKA), Aurora kinase B (AURKB), and Aurora kinase C (AURKC). AURK expression varies throughout the cell cycle and its activity is regulated through a combination of phosphorylation, localization, and cofactors ^14,15^. Dephosphorylation of AURKs plays an important role in both activity and degradation. Specifically, removal of a phosphate group at Ser51 by the PP2A complex initiates AURK degradation and is associated with mitotic exit^14^. While AURKs play key roles in cell cycle regulation, a considerable number of non-mitotic functions of AURKs are also being studied, especially regarding DNA damage response ^15^. Thus far, Vif-dependent AURK activation has mainly been studied in relation to cell cycle arrest because prolonged AURKA and AURKB activation is associated with failure of mitotic exit ^1^. With this study, we aimed to investigate another possible and largely unexamined outcome of Vif-induced AURK dysregulation: regulation of cell-cell fusion.

HIV-1 transmission occurs largely through cell-free virion release or through cell-to-cell transmission. Both cell-free virion release and cell-to-cell transmission require the assembly and egress of viral particles. However, in cell-to-cell transmission, viral budding and assembly are polarized at a site of cell-cell contact termed the virological synapse (VS), which allows for highly efficient transfer of virus to the target T cell ^16–22^. VS formation requires the fusogenic viral glycoprotein Env ^18^, which can induce membrane fusion at physiological pH. In most cases, both *in vivo* and *in vitro*, VS formation results in efficient transmission of viral particles and no fusion between producer and target cells ^23^. Alternatively, VS formation can result in cell-cell fusion, i.e. the fusion of producer and target cells, giving rise to multinucleated cells called syncytia. HIV-1-induced, purely T cell-based syncytia have been documented *in vivo* ^23–29^ and are typically small (2-5 nuclei) and infrequent (∼10-20% of infected T cells) ^27^. Since this low level of fusion occurs and appears to be carefully controlled, cell-cell fusion regulation is likely an important process for successful spread of the virus.

Both viral and host mechanisms operate to inhibit excess cell-cell fusion at the VS. In the absence of Gag, Env is downregulated from the surface through rapid endocytosis, thus preventing cell-cell fusion ^30^. In addition, immature Gag traps Env in a poorly fusogenic state through interaction with Env’s cytoplasmic tail, inhibiting Env-induced fusion until maturation ^31–33^. Tetraspanins ^34–39^ and the tetraspanin partner EWI-2 ^40^ accumulate at the HIV-1 VS and prevent excess cell-cell fusion. Another tetraspanin partner, the Ezrin-radixin-moesin (ERM) protein Ezrin, more specifically its Thr567-phosphorylated form (p-Ezrin-T567), also accumulates at the VS and inhibits excess HIV-1-induced cell-cell fusion ^41^. Ezrin phosphorylation at Thr567 occurs through by ROCK, a cellular kinase and downstream effector of the GTPase RhoA ^42,43^. It is still unclear how Ezrin is phosphorylated at the VS, and it likely involves multiple RhoA/ROCK-related proteins.

One possible connection that explains fusion-inhibitory Ezrin-T567 phosphorylation at the VS is that the cytoplasmic domain of Env (Env CTD) has been shown to interact with p115-RhoGEF (ARHGEF1) ^44^. This interaction may provide a bridge between viral and host proteins, facilitating localized activation Ezrin. However, there is also a plausible connection between AURK activity and p-Ezrin-T567: 14-3-3 binds the kinesin component of the centralspindlin complex ^45^ to prevent its oligomerization and plasma membrane localization ^46^. During cleavage furrow formation in cytokinesis, active AURKB relieves 14-3-3-mediated inhibition of centralspindlin ^45,46^, resulting in activation of the plasma membrane-associated RhoGEF ECT2 (ARHGEF31) which subsequenty activates RhoA ^46^ and induces ROCK-mediated Ezrin-T567 phosphorylation.

Previous investigations in our lab revealed a novel relationship between Vif and cell-cell fusion: we found that Vif-null virus exhibited significantly higher levels of cell-cell fusion compared to Vif-proficient virus in several complementary assays ^47^. Separately, another study found an unexpected connection between AURKB activity and cell-cell fusion ^48^. Inhibition of AURKB in HIV-1-infected cells caused an increase in Env-mediated cell-cell fusion, specifically through interaction with Env cytoplasmic tail domain ^48^. Taken together, these findings point to a highly plausible connection between Vif, AURK activity, and cell-cell fusion regulation in HIV-1-infected T cells.

The goal of this study was to elucidate how these factors are functionally connected (Figure 1). We hypothesize that, through degradation of PPP2R5A-E, Vif helps limit the rate of Env-mediated cell-cell fusion at the virological synapse by increasing AURK pathway activity (Figure 2). Using split Nano Luciferase (split-NLuc)-based HIV-1-induced cell-cell fusion assays along with a small panel of AURK inhibitors, we evaluated the impact of Vif on cell-cell fusion. Potential AURK downstream targets and proteins involved with fusion regulation were evaluated with Western blotting. In support of our hypothesis, we found that Vif-mediated cell-cell fusion regulation was dependent on AURK activity, and that Ezrin-T567 phosphorylation was elevated in the presence of Vif.

**Figure 1.**
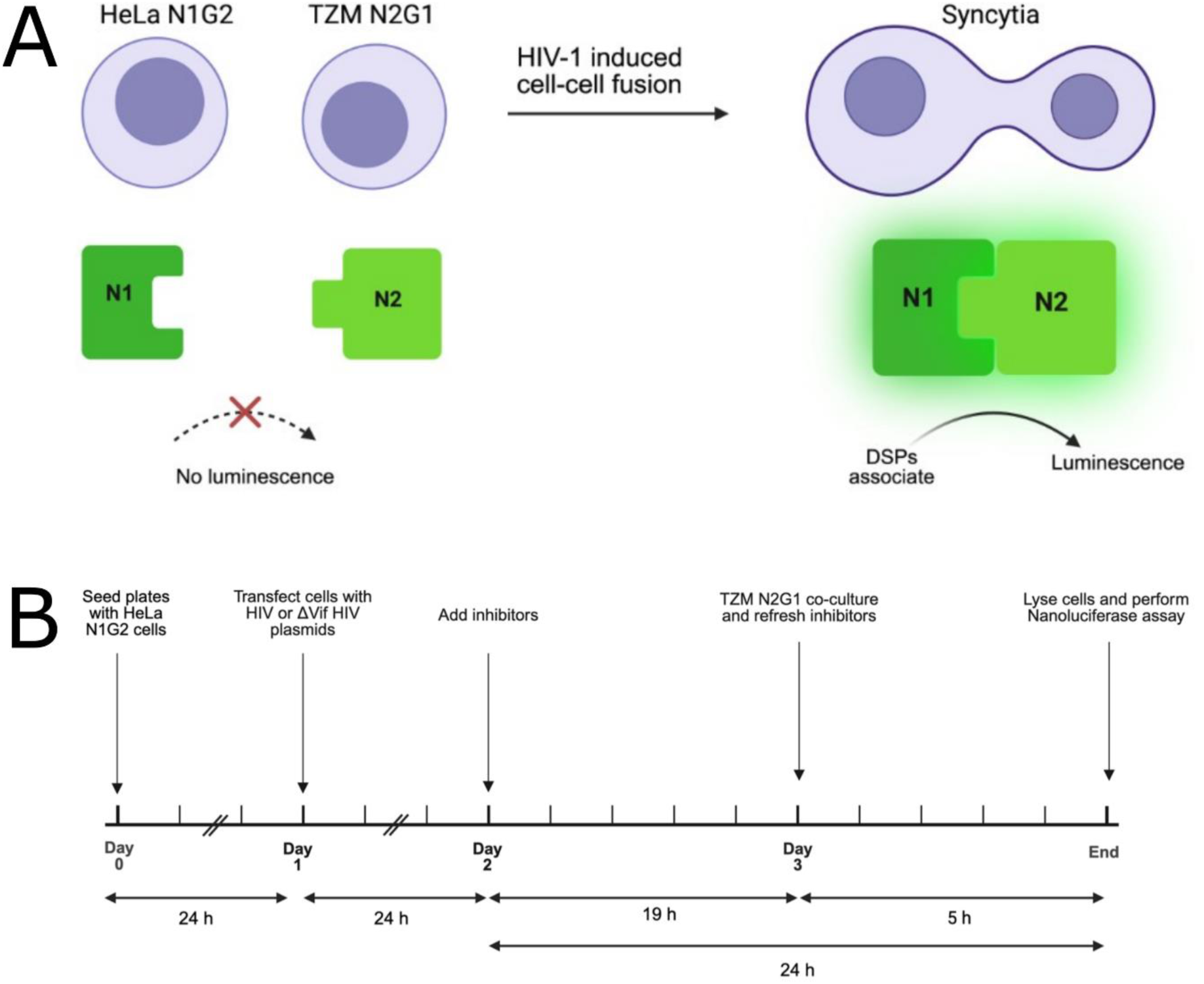
Split-NLuc cell-cell fusion assay. **(A)** Assay schematic. HeLa and TZM cells were transduced and selected to constitutively express NLuc-GFP-based dual split protein (DSP) system. Upon cell-cell fusion, the split reporter components combine. Luminescence signal upon addition of NanoGlo reagent (Promega) to cell lysates corresponds to cell-cell fusion activity (syncytium formation). **(B)** Experiment timeline. HeLa-N1G2 cells are plated on day zero, transfected with proviral plasmid 24 h later, and drugs are added 24 h after transfection in fresh media. The next day, TZM-bl-N2G1 cells are added for a 5 h coculture, followed by cell lysis and NLuc activity assessment by NanoGlo assay (Promega).

**Figure 2.**
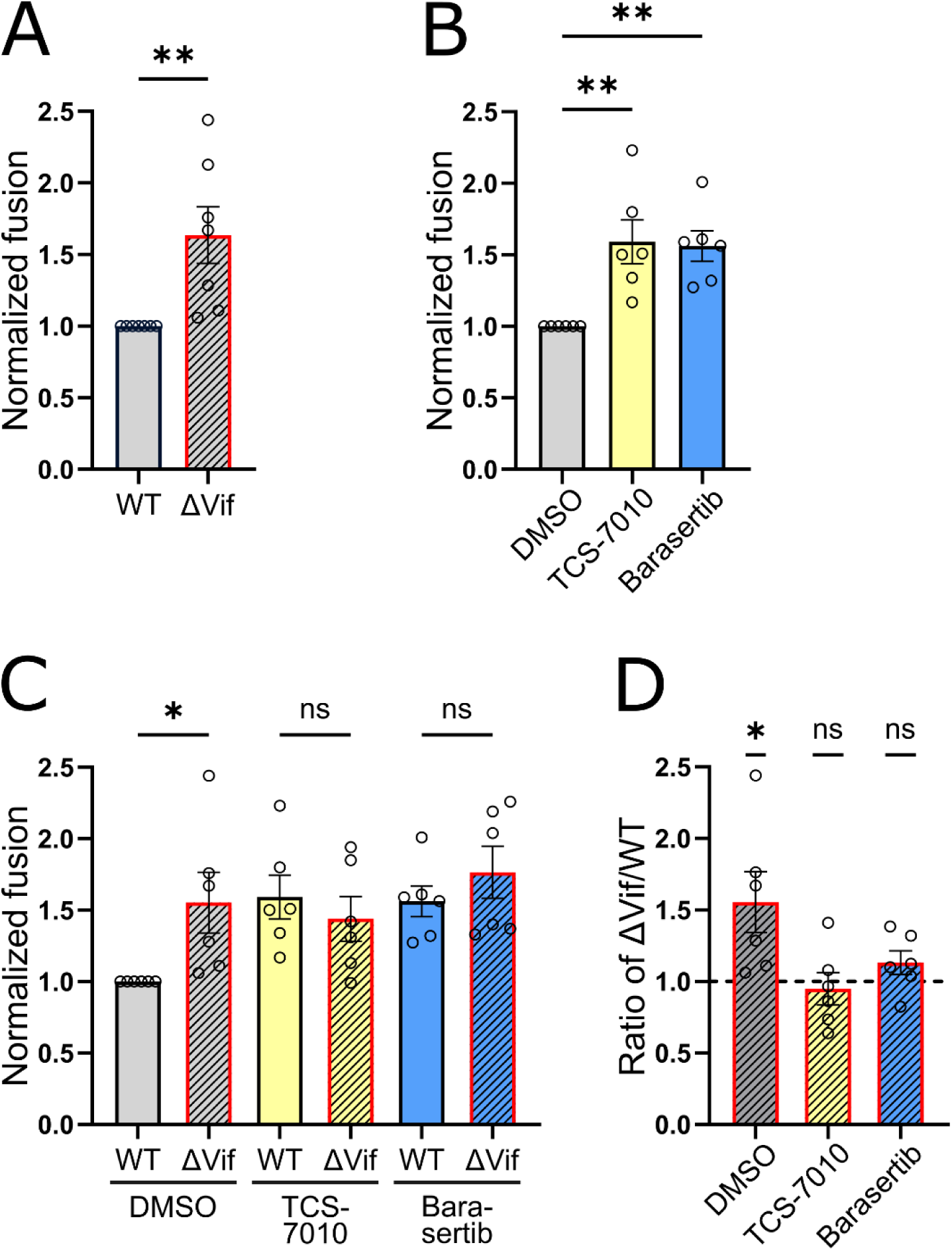
Vif and AURKs operate through a common cell-cell fusion regulation mechanism. Cell-cell fusion between HIV-1-transfected HeLa-N1G2 cells and TZM-bl-N2G1 cells was evaluated using split-NLuc assays. **(A)** Wild-type (WT) and Vif-deleted (ΔVif) conditions, normalized to WT. Unpaired t-test, **: p ≤ 0.01. **(B)** Cells treated with vehicle control (DMSO), AURKA inhibitor (TCS-7010, 5 μM), or AURKB inhibitor (Barasertib, 10 μM), with WT HIV-1 transfection only, normalized to DMSO. Kruskal-Wallis test (vs. DMSO) with Dunn’s correction for multiple comparisons, **: p ≤ 0.01. **(C)** Cells treated with DMSO, TCS-7010 (5 μM), or Barasertib (10 μM), with WT or ΔVif transfection, normalized to WT + DMSO. One-way ANOVA comparing WT to ΔVif within each drug treatment, *: p ≤ 0.05. **(D)** The data in (C) were reanalyzed by dividing the ΔVif values by the WT values. One sample t test comparing each dataset to the theoretical value of 1 (= no Vif-dependent effect), *: p ≤ 0.05. **(A-D)** Each datapoint shown (open circles) represents one biologically-independent replicate, each representing the average of 5 technical replicates after elimination of the highest and lowest value among those replicates. Bars represent the mean ± SEM of biological replicates.

## 2. Materials and Methods

### 2.1. Cell Lines and Cell Culture

HeLa and TZM-bl cells (which are HeLa cells rendered susceptible to HIV infection through constitutive expression of CD4 and CCR5) were obtained through BEI Resources, NIAID, NIH: HeLa Cells, HRP-153; TZM-bl Cells, HRP-8129. Both cell lines were maintained at 37°C in 5% CO2/95% humidity in Dulbecco’s Modified Eagle’s Medium (DMEM; Corning #10-017-CV) with 10% fetal bovine serum (FBS; Cytiva #SH30910.03), with the addition of 0.25 µg/mL puromycin (Gibco #A1113803) for stable selection of the transgenes described below. For passaging cells, 1X TrypLE Express Enzyme (Gibco #12604021) was used as a dissociation reagent. All experiments were conducted in media supplemented with 10% FBS and without any antibiotics. HeLa and TZM-bl cells were stably transduced with pLX301-based lentiviral vectors for expression of split-NLuc constructs: HeLa cells with N1G2 (the C-terminal portion of GFP fused to the N-terminal portion of Nano Luficerase), and TZM-bl cells with N2G1 (the N-terminal portion of GFP fused to the C-terminal portion of Nano Luciferase), adapted from the GFP-Rluc8 dual split protein (DSP) system published by the Matsuda group ^49,50^. Upon fusion between HeLa-N1G2 with TZM-bl-N2G1 cells, these two constructs irreversibly combine, resulting in both Nano Luciferase activity and GFP fluorescence. Cells were routinely checked for mycoplasma by PCR; no positivity was detected at any time.

### 2.2. Plasmid DNA

HIV IIIB proviral plasmids (WT and ΔVif) were originally provided by Reuben Harris. HIV ΔVif is identical to the WT HIV provirus, but contains an irreparable 230 bp deletion in the *vif* gene ^47^. Plasmid stocks were generated by transforming chemically competent STBL3 *E. coli* cells by standard heat shock procedure. Transformed bacteria were plated on LB agar containing 100 µg/mL carbenicillin, grown at room temperature for 48 h, and colonies were picked and shaken for 48 h at 30°C in 5 mL LB/carbenicillin (100 µg/mL) broths. Plasmid DNA was isolated from the broths and run on a 0.7% agarose gel to screen for absence of plasmid recombination. Successful transformants were further expanded in 250 mL LB/carbenicillin broths at 30°C, and plasmid DNA was purified using the E.Z.N.A. Plasmid Maxi kit (Omega Bio-tek #D6922-04) before being screened for absence of recombination again. Proteinase K was added at the same time as lysis solution during all plasmid preparation due to expression of *endA* in STBL3 *E. coli*.

### 2.3. Aurora Kinase Inhibitors and Cell Viability Assays

AURK-targeting chemical inhibitors (Table 1) with high specificity towards either AURKA or AURKB were selected for use in cell-cell fusion assays. Chemical inhibitors were reconstituted from powder in DMSO at 10 mM and stored in single-use aliquots at -80°C to avoid repeated freeze-thaw cycles. These chemicals were used at working concentrations selected after testing for cell viability at concentrations bracketing published IC50 values for each. Cell viability was measured using a commercial MTS assay (CellTiter 96 AQueous Non-Radioactive Cell Proliferation Assay, Promega #G5421). HeLa cells were seeded in 96-well plates at 1.7 x 10^4^ cells per well in 100 µL of complete growth medium and incubated for 48 h. Cells were then treated with a range of concentrations of each drug or vehicle control for 24 h, before addition of 5 µL of MTS reagent per well. Plates were incubated for 45 min at 37°C in before measuring absorbance at 490 nm by plate reader. Wells containing only media and MTS reagent without cells were used as blanks. Each treatment was evaluated in five technical replicates. Cell viability was calculated as a percentage of vehicle control.

**Table 1.** Kinase inhibitors used in cell-cell fusion assays.

| **Drug Name** | **Target** | **Catalog No.** | **CAS Number** | **Source** |
| --- | --- | --- | --- | --- |
| TCS-7010 | AURKA | S1451 | 1158838-45-9 | Selleck Chemicals |
| Barasertib | AURKB | 50-780-2 | 722544-51-6 | Selleck Chemicals |

### 2.5. Cell-Cell Fusion Assays

Split-NLuc assays (Figure 1) were used to quantify cell-cell fusion. HeLa-N1G2 cells were seeded on 96-well plates at 1.7 x 10^4^ cells per well, and 24 h later transfected with HIV-1 proviral plasmids (IIIB WT or Δvif; 170 ng of DNA per well) using FuGENE6 (Promega #E2692). 24 h after transfection, media were refreshed with drug or vehicle-containing media, and another 19 h later, drug/vehicle-containing media were refreshed and 8 x 10^4^ TZM-bl-N2G1 cells (lifted using PBS/15 mM EDTA and washed) were added per well. After 5 h, supernatants were removed and wells were lysed with 75 μL lysis buffer per well (5x Reporter Lysis Buffer; Promega #E3971, 100x Mammalian Protease Inhibitor Cocktail; ApexBio # K1007, freshly diluted to 1x in sterile double-distilled water). Plates were shaken at high speed for 15 min. 50 µL of each cell lysate was combined with 50 µL of NanoGlo reagent (Promega #N1120) in a white 96-well plate (Thermo Scientific #7571). Wells containing lysis buffer and NanoGlo were used for background subtraction. Luminescence was measured by plate reader after a 3 min incubation at room temperature in the dark. All treatments were evaluated with five technical replicates per assay, and a set of T-20-treated IIIB WT-transfected cell-cell fusion negative controls were run within each assay.

### 2.6. Western Blots

HeLa-N1G2 cells were plated in 12-well plates at 1.8 x 10^5^ cells per well and 24 h later transfected with HIV-1 IIIB (WT or ΔVif) proviral plasmid using FuGENE6 as described above. 48 h later, cells were washed and lifted with PBS/15 mM EDTA, pelleted, and lysed for 10 min with ice-cold 1x RIPA lysis buffer (5X RIPA Lysis Buffer; Thermo Scientific #J62524AD, 100x Mammalian Protease Inhibitor Cocktail; ApexBio #K1007, 100x Halt Phosphatase Inhibitor Single-Use Cocktail; Thermo Scientific #78420, freshly diluted to 1x in sterile double-distilled water). Lysates were centrifuged at 20,000 rcf for 5 min at 4°C and supernatants were transferred into fresh tubes. Protein was quantified using the Pierce Detergent Compatible Bradford Assay Kit (Thermo Scientific #23246). Samples were mixed at 3:1 ratio with 4X Reducing Laemmli sample buffer (Thermo Scientific #J60015AC) and boiled at 95°C for 5 min, and 15 µg of protein was loaded per lane alongside PageRuler Prestained Protein Ladder (Thermo Scientific #26616). Samples were electrophoresed on homemade 8% resolving/4% stacking polyacrylamide gels in 1X SDS-PAGE running buffer and transferred to PVDF membrane (Millipore #IPVH00010). Membranes were blocked in blocking buffer (1X Tris Buffered Saline; TBS, containing 0.1% Tween 20; TBST, with 5% non-fat milk powder) for 1 h at 4°C. Membranes that would be blotted for phosphorylated proteins were instead blocked in 1X TBST containing 5% BSA for up to 4 h at 4°C. Primary antibodies were incubated overnight shaking at 4°C in either TBST-milk or TBST-BSA buffer. Antibodies and dilutions used are listed in Table 2. Membranes were washed in 1X TBST and horseradish peroxidase (HRP)-conjugated secondary antibodies (Table 2) were incubated in TBST-milk buffer at room temperature for 1 h. Membranes were washed in 1X TBST, signal was developed with SuperSignal West Femto Maximum Sensitivity Substrate (Thermo Scientific #34095), and blots were imaged on a ChemiDoc imager (Bio-Rad).

**Table 2.** Antibodies used in Western blotting.

| **Antibody** | **Catalog No.** | **Source** | **Dilution** |
| --- | --- | --- | --- |
| Ezrin | 3145 | Cell Signaling Technology | 1:1,000 |
| Phospho-Ezrin (Thr567)/Radixin (Thr564)/Moesin (Thr558) (48G2) | 3726 | Cell Signaling Technology | 1:250 |
| Mouse anti-HIV-1 p24 (AG3.0) | 3537 | NIH HIV Reagent Program | 1:1,000 |
| GAPDH Monoclonal Antibody (6C5) | AM4300 | Invitrogen | 1:100,000 |
| Histone H3 | PA5-16183 | Invitrogen | 1:40,000 |
| Phospho-Histone H3 (Ser10) | 701258 | Invitrogen | 1:500 |
| AURKA | 14475 | Cell Signaling Technology | 1:1,000 |
| AURKB | 3094 | Cell Signaling Technology | 1:1,000 |
| Phospho-Aurora A (Thr288)/Aurora B (Thr232)/Aurora C (Thr198) | 2914 | Cell Signaling Technology | 1:500 |
| Donkey anti-Rabbit HRP | 711-035-152 | Jackson ImmunoResearch | 1:1,000 |
| Donkey anti-Mouse HRP | 715-035-150 | Jackson ImmunoResearch | 1:1,000 |

## 3. Results

### 3.1. Vif and AURKs regulate HIV-1-induced cell-cell fusion through a common pathway

HIV-1 Vif ^47^ and AURKs ^48^ have been independently implicated as inhibitors of HIV-1-induced cell-cell fusion. We first sought to recapitulate those earlier findings in a newly-developed highly quantitative HeLa/TZM-bl cell-based fusion assay utilizing split-NLuc (Figure 1). Our results (Figure 2) closely recapitulated the earlier report ^47^ that deletion of Vif leads to a significant increase in cell-cell fusion (Figure 2A). Furthermore, we confirmed ^48^ that AURKB inhibits cell-cell fusion (Figure 2B; Barasertib treatment at 10 μM). However, in contrast with that previous study, we found that AURKA acts similarly to AURKB (Figure 2B; TCS-7010 treatment at 5 μM), i.e. inhibition of either AURKA or AURKB led to a significant increase in cell-cell fusion activity. We confirmed that these inhibitors did not impact cell viability at the concentrations used in cell-cell fusion assays (data not shown). Empty vector-transfected cells and HIV-1-transfected cells treated with an Env-specific fusion inhibitor (T-20) yielded negligible signal (data not shown), confirming the specificity of our split-NLuc-based fusion assay. Together, these results laid the necessary foundation which allowed us to go on to test whether Vif and AURKs function as part of the same fusion-regulating pathway.

Vif manipulates the activity of the protein phosphatase PP2A, which has downstream impacts on AURK signaling ^1,10^. We sought to functionally evaluate the relationship between Vif and AURKs as cell-cell fusion inhibitors by measuring cell-cell fusion activity while simultaneously deleting Vif and inhibiting AURKs. When normalizing values to the WT/vehicle control condition, we observed that all treatments increased cell-cell fusion (Figure 2C), and statistical comparison between WT and ΔVif conditions within each drug treatment showed that only the vehicle control exhibited a significant Vif-dependent effect. In other words, AURK inhibition occludes the effect of Vif deletion.

To analyze these same data in a different way, we divided the ΔVif values by the WT values within each drug treatment, and compared the resulting dataset to the theoretical value of 1 (Figure 2D). Any values significantly higher than 1 would indicate that Vif deletion was causing an increase in cell-cell fusion activity (as already shown in Figure 2A). In agreement with the first analysis (Figure 2C), only vehicle-treated cells exhibited a value significantly higher than 1, indicating Vif-dependent fusion regulation (Figure 2D). Both TCS-7010- and Barasertib-treated cells exhibited values indistinguishable from the theoretical value of 1, indicating no effect of Vif on fusion in those conditions where AURKA/B were inhibited. These two analyses indicated that AURK inhibition and Vif deletion exhibit a non-additive interaction, which is consistent with the hypothesis that Vif and AURKs operate within the same cell-cell fusion regulation pathway.

### 3.2. Vif alters the Phosphorylation Status of Host Proteins Involved in Regulating Cell-Cell Fusion

Upon establishing that Vif-mediated inhibition of HIV-1-induced cell-cell fusion operates through AURK signaling, we asked whether any other known fusion regulators are possible targets of AURKs and whether their phosphorylation status is altered when Vif is deleted. One such candidate fusion regulator is Ezrin; we previously showed that p-Ezrin-T567 accumulates at the VS and inhibits cell-cell fusion ^41^, and Ezrin phosphorylation is likely to be influenced by AURKs through multiple pathways, as reviewed above in Introduction. We thus hypothesized that Vif influences the phosphorylation status of Ezrin, specifically, it increases the level of p-Ezrin-T567, which in turn acts as a fusion inhibitor ^41^.

To address whether Ezrin phosphorylation is controlled by Vif, we transfected HeLa cells with WT or ΔVif HIV-1 proviral plasmids, or empty vector, then lysed cells and performed Western blotting for viral and host proteins/phosphoproteins of interest. Strikingly, we observed that, while total Ezrin levels were unaffected by Vif deletion, p-Ezrin-T567 was virtually undetectable in ΔVif conditions (Figure 3A). Note that, while the p-ERM antibody used to detect p-Ezrin-T567 is cross-reactive with a highly similar phosphosite on the other two ERM proteins (Radixin and Moesin), the complete loss of any p-ERM signal in ΔVif conditions, and because we have previously documented that Ezrin is the functionally relevant ERM protein in HIV-1-infected cells ^41^, we can reasonably conclude that Ezrin-T567 phosphorylation is lost when Vif is deleted.

**Figure 3.**
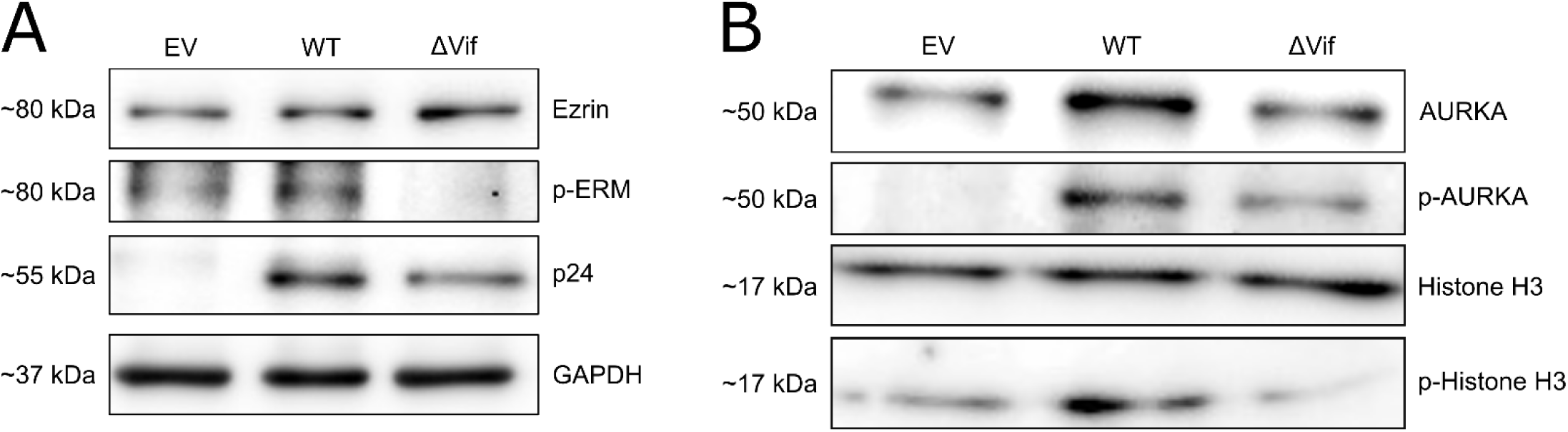
AURK signaling and Ezrin phosphorylation are altered in the absence of Vif. HeLa-N1G2 cells were transfected with wild-type (WT) or Vif-deleted (ΔVif) HIV-1 IIIB proviral plasmid, or empty vector (EV). Cells were then lysed and proteins were analysed by Western blotting. **(A)** Ezrin-T567 phosphorylation is lost when Vif is deleted. Phospho-ERM (pERM) includes the phosphorylated forms of all three ERM proteins, but the total Ezrin antibody does not cross-react with other ERM proteins. **(B)** AURKA and its downstream target Histone H3 are hyperphosphorylated in the presence of Vif (WT) and dephosphorylated in the absence of Vif (ΔVif).

To confirm that AURKs are themselves maintained at a hyperphosphorylated state by Vif (due to PP2A downregulation) ^1,10^, we also performed Western blotting for total and phosphorylated AURKA and Histone H3 (one of the direct phosphorylation targets of AURKA). As expected, and in agreement with several previous reports, we found that AURKA and Histone H3 were hyperphosphorylated in cells transfected with WT HIV-1 compared to empty vector-transfected cells, and that this effect was dramatically reversed when Vif was deleted (Figure 3B). Taken together, our results support a cell-cell fusion-inhibitory mechanism involving Vif, AURKs, and Ezrin.

## 4. Discussion

To secure efficient virus spread via cell-to-cell transmission, HIV-1 has evolved to inhibit excessive Env-induced cell-cell fusion. Because Env is fusogenic at physiological pH, its fusogenicity at the HIV-1 VS is regulated in other ways, including its rapid retrieval from the surface of the cell ^30^, and Gag-mediated restriction of Env mobility through an interaction with Env’s cytoplasmic tail ^31–33^. Also, Gag recruits tetraspanins to the VS ^34^ where they ^38,39^, together with partner proteins EWI-2 ^40^ and Ezrin ^41^ help inhibit excess fusion. Ezrin must be in its phosphorylated/active state (p-Ezrin-T567) to interact with tetraspanins and the cytoskeleton and thus inhibit membrane fusion at the HIV-1 VS. While it is possible that HIV-1 passively depends on Ezrin phosphorylation to occur to mediate this fusion regulation, HIV-1 infection could instead actively promote Ezrin phosphorylation to ensure that this mechanism is operational. With this report, we now demonstrate that this is indeed the case: we find that HIV-1 Vif orchestrates the hyperactivation of AURKs, which results in the downstream phosphorylation of Ezrin-T567 to secure excess cell-cell fusion is prevented. Deletion of Vif completely reverses this regulatory regime and results in uncontrolled cell-cell fusion.

This investigation was prompted in part by a recent report from the Sherer lab that AURKB activity was involved in regulating cell-cell fusion ^48^, combined with prior research from our lab that identified an unknown role for Vif in fusion regulation ^47^, and the initial finding that AURK signaling is dysregulated by Vif in HIV-1-infected cells ^10^. While our results (Figure 2B) are in agreement with the Sherer lab that Barasertib, an AURKB inhibitor, causes an increase in cell-cell fusion ^48^, our finding that TCS-7010, an AURKA inhibitor, also does the same is discrepant with the Sherer lab’s report. The cell-cell fusion assays used by the Sherer lab involved HIV-1-expressing Jurkat T cells as the producer cells, whereas in the present study we used HeLa cells transfected with HIV-1. In both cases, target cells were TZM-bl cells. Another key difference was the timeline of the experiment; in the Sherer lab report, pretreatment with inhibitors was done for 2 h (whereas we pretreated for 19 h), followed by 2 h of coculture (whereas we cocultured for 5 h), after which producer cells were washed off and cells were cultured for another 48 h to allow reporter expression. Because we used a split-NLuc system which requires no additional incubation time to develop signal as the reassembled NLuc enzyme is immediately active, we were able to assay for cell-cell fusion activity directly after the end of the 5 h coculture period. The concentrations we used for both drugs (TCS-7010 and Barasertib) were half those used by the Sherer lab. Altogether, we think it is likely that the fusion increase we observed upon treatment with the AURKA inhibitor TCS-7010 would have also been observed by the Sherer lab if they had used a longer pretreatment period, and/or if they did not need to culture cells for another 48 h after the coculture period. Our findings are generally in agreement that AURK signaling regulates HIV-1-induced cell-cell fusion.

Sub-cellular localization of AURKs may prove to be important to their role in fusion regulation. Both AURKA and AURKB activity involve phosphorylation of structures within and around centrosomes. Increased activity of AURKs and phosphorylation of targets proximal to the VS directly aligns with the cytoskeletal polarization seen in the producer cell during cell-cell transmission at the VS ^51^. In HIV-1-infected T cells, the MTOC is reoriented to the site of cell-cell contact, positioning the centrosomes directly beneath the virological synapse ^22,51^. The association of activated AURKA and AURKB with the MTOC and centrosome-associated proteins likely allows their polarized localization at the VS to locally orchestrate the necessary conditions to promote Ezrin-T567 phosphorylation. More specifically, we predict that AURKA/B locally de-repress and stabilize the centralspindlin complex, which subsequently activates ECT2, and that in turn activates ROCK which phosphorylates Ezrin-T567 ^45,46,52^. Phosphorylated Ezrin is then able to simultaneously interact with the cytoplasmic domains of transmembrane proteins (such as the tetraspanin CD81, which also localizes to the HIV-1 VS) and the actin cytoskeleton, resulting in stiffening of the membrane milieu surrounding the VS, and thus physically preventing cell-cell fusion.

While our prior and ongoing work has mostly focused on HIV-1-induced small T cell syncytia and their potential role as efficient, evasive, and mobile virus spreaders ^27^, the present study leads us to ask whether pharmacologically dysregulating fusion may prove useful in a therapeutic context. Uncontrolled fusion leading to an accumulation of large syncytia may accelerate depletion of the virus reservoir, either through direct cytopathic effects, or even through increased immune detection and killing; in a preprint (Girard *et al*., 2026, bioRxiv) submitted simultaneously with this one, we present evidence that syncytia are significantly better targeted and killed by natural killer (NK) cells than infected mononucleated cells. This suggests that AURK inhibition could be an effective antiviral strategy, including during “shock-and-kill” strategies for HIV-1 cure. Other kinase inhibitors may also prove to be useful in this regard, as certain ones have been shown to increase cell-cell fusion for other syncytium-forming viruses, including as measles and SARS-CoV-2 ^53,54^.

Cell-cell fusion is a major challenge for a virus such as HIV-1, which spreads efficiently via cell-to-cell transmission. Syncytium formation is the opposite of virus transmission; transmission amplifies the number of infected cells, while syncytium formation merely grows the size of one infected cell. However, a tightly regulated, low level of cell-cell fusion might be advantageous for viral spread through formation of small motile syncytia that participate in virus transmission and perhaps possess unique properties which overall promote spread. Regulation of cell-cell fusion is therefore essential for viral homeostasis. This study reveals key mechanistic details of how HIV-1 Vif usurps AURK signaling to help secure the integrity of the VS during cell-to-cell transmission.

## 5. Acknowledgments

## 6. Author Contributions

SJT: investigation, data analysis, manuscript authorship and editing. BL: investigation, data analysis. JPG: investigation. EM: investigation. MKC: investigation. MT: supervision, funding acquisition, data analysis, manuscript editing. MS: concept origination, supervision, data analysis, manuscript authorship and editing. Large language model artificial intelligence (LLM-AI) was not utilized anywhere in this work.

## References

1 Salamango, D. J., Ikeda, T., Moghadasi, S. A., Wang, J., McCann, J. L., Serebrenik, A. A., Ebrahimi, D., Jarvis, M. C., Brown, W. L. & Harris, R. S. HIV-1 Vif Triggers Cell Cycle Arrest by Degrading Cellular PPP2R5 Phospho-regulators. Cell Rep 29, 1057–1065 e1054 (2019). PubMed PMC6903395

2 Harris, R. S., Hultquist, J. F. & Evans, D. T. The restriction factors of human immunodeficiency virus. J. Biol. Chem. 287, 40875–40883 (2012). PubMed PMC3510791

3 Land, A. M., Wang, J., Law, E. K., Aberle, R., Kirmaier, A., Krupp, A., Johnson, W. E. & Harris, R. S. Degradation of the cancer genomic DNA deaminase APOBEC3B by SIV Vif. Oncotarget 6, 39969–39979 (2015). PubMed PMC4741873

4 Zhao, R. Y. & Elder, R. T. Viral infections and cell cycle G2/M regulation. Cell Res. 15, 143–149 (2005). PubMed 15780175

5 Sakai, K., Dimas, J. & Lenardo, M. J. The Vif and Vpr accessory proteins independently cause HIV-1-induced T cell cytopathicity and cell cycle arrest. Proc Natl Acad Sci U S A 103, 3369–3374 (2006). PubMed PMC1413893

6 DeHart, J. L., Bosque, A., Harris, R. S. & Planelles, V. Human immunodeficiency virus type 1 Vif induces cell cycle delay via recruitment of the same E3 ubiquitin ligase complex that targets APOBEC3 proteins for degradation. J. Virol. 82, 9265–9272 (2008). PubMed PMC2546900

7 Salamango, D. J. & Harris, R. S. Demystifying Cell Cycle Arrest by HIV-1 Vif. Trends Microbiol. 29, 381–384 (2021). PubMed PMC8416294

8 Salamango, D. J. & Harris, R. S. Dual Functionality of HIV-1 Vif in APOBEC3 Counteraction and Cell Cycle Arrest. Front Microbiol 11, 622012 (2020). PubMed PMC7835321

9 Bandini, M., Ghone, D., Evans, E. L., 3rd, Suzuki, A. & Sherer, N. M. HIV-1 Vif and Vpr cooperatively modulate the cell cycle to maximize per-cell virion production. Proc Natl Acad Sci U S A 122, e2511502122 (2025). PubMed PMC12626010

10 Greenwood, E. J., Matheson, N. J., Wals, K., van den Boomen, D. J., Antrobus, R., Williamson, J. C. & Lehner, P. J. Temporal proteomic analysis of HIV infection reveals remodelling of the host phosphoproteome by lentiviral Vif variants. Elife 5 (2016). PubMed PMC5085607

11 Hu, Y., Delviks-Frankenberry, K. A., Wu, C., Arizaga, F., Pathak, V. K. & Xiong, Y. Structural insights into PPP2R5A degradation by HIV-1 Vif. Nat. Struct. Mol. Biol. 31, 1492–1501 (2024). PubMed PMC12221852

12 Salamango, D. J., McCann, J. L., Demir, O., Becker, J. T., Wang, J., Lingappa, J. R., Temiz, N. A., Brown, W. L., Amaro, R. E. & Harris, R. S. Functional and Structural Insights into a Vif/PPP2R5 Complex Elucidated Using Patient HIV-1 Isolates and Computational Modeling. J. Virol. 94 (2020). PubMed PMC7565612

13 Johnson, J. R., Crosby, D. C., Hultquist, J. F., Kurland, A. P., Adhikary, P., Li, D., Marlett, J., Swann, J., Huttenhain, R., Verschueren, E., Johnson, T. L., Newton, B. W., Shales, M., Simon, V. A., Beltrao, P., Frankel, A. D., Marson, A., Cox, J. S., Fregoso, O. I., Young, J. A. T. & Krogan, N. J. Global post-translational modification profiling of HIV-1-infected cells reveals mechanisms of host cellular pathway remodeling. Cell Rep 39, 110690 (2022). PubMed PMC9429972

14 Willems, E., Dedobbeleer, M., Digregorio, M., Lombard, A., Lumapat, P. N. & Rogister, B. The functional diversity of Aurora kinases: a comprehensive review. Cell Div 13, 7 (2018). PubMed PMC6146527

15 Ma, H. T. & Poon, R. Y. C. Aurora kinases and DNA damage response. Mutat. Res. 821, 111716 (2020). PubMed 32738522

16 Phillips, D. M. The role of cell-to-cell transmission in HIV infection. AIDS 8, 719–731 (1994). PubMed 8086128

17 Johnson, D. C. & Huber, M. T. Directed egress of animal viruses promotes cell-to-cell spread. J. Virol. 76, 1–8 (2002). PubMed PMC135733

18 Jolly, C., Kashefi, K., Hollinshead, M. & Sattentau, Q. J. HIV-1 cell to cell transfer across an Env-induced, actin-dependent synapse. J. Exp. Med. 199, 283–293 (2004). PubMed PMC2211771

19 Jimenez-Baranda, S., Gomez-Mouton, C., Rojas, A., Martinez-Prats, L., Mira, E., Ana Lacalle, R., Valencia, A., Dimitrov, D. S., Viola, A., Delgado, R., Martinez, A. C. & Manes, S. Filamin-A regulates actin-dependent clustering of HIV receptors. Nat. Cell Biol. 9, 838–846 (2007). PubMed 17572668

20 Chen, P., Hubner, W., Spinelli, M. A. & Chen, B. K. Predominant mode of human immunodeficiency virus transfer between T cells is mediated by sustained Env-dependent neutralization-resistant virological synapses. J. Virol. 81, 12582–12595 (2007). PubMed PMC2169007

21 Hubner, W., McNerney, G. P., Chen, P., Dale, B. M., Gordon, R. E., Chuang, F. Y., Li, X. D., Asmuth, D. M., Huser, T. & Chen, B. K. Quantitative 3D video microscopy of HIV transfer across T cell virological synapses. Science 323, 1743–1747 (2009). PubMed PMC2756521

22 Starling, S. & Jolly, C. LFA-1 Engagement Triggers T Cell Polarization at the HIV-1 Virological Synapse. J. Virol. 90, 9841–9854 (2016). PubMed PMC5068534

23 Compton, A. A. & Schwartz, O. They Might Be Giants: Does Syncytium Formation Sink or Spread HIV Infection? PLoS Pathog 13, e1006099 (2017). PubMed PMC5289631

24 Orenstein, J. M. In vivo cytolysis and fusion of human immunodeficiency virus type 1-infected lymphocytes in lymphoid tissue. J. Infect. Dis. 182, 338–342 (2000). PubMed 10882620

25 Orenstein, J. M. HIV expression in surgical specimens. AIDS Res. Hum. Retroviruses 24, 947–955 (2008). PubMed 18671477

26 Murooka, T. T., Deruaz, M., Marangoni, F., Vrbanac, V. D., Seung, E., von Andrian, U. H., Tager, A. M., Luster, A. D. & Mempel, T. R. HIV-infected T cells are migratory vehicles for viral dissemination. Nature 490, 283–287 (2012). PubMed PMC3470742

27 Symeonides, M., Murooka, T. T., Bellfy, L. N., Roy, N. H., Mempel, T. R. & Thali, M. HIV-1-Induced Small T Cell Syncytia Can Transfer Virus Particles to Target Cells through Transient Contacts. Viruses 7, 6590–6603 (2015). PubMed PMC4690882

28 Law, K. M., Komarova, N. L., Yewdall, A. W., Lee, R. K., Herrera, O. L., Wodarz, D. & Chen, B. K. In Vivo HIV-1 Cell-to-Cell Transmission Promotes Multicopy Micro-compartmentalized Infection. Cell Rep 15, 2771–2783 (2016). PubMed 27292632

29 Ventura, J. D., Beloor, J., Allen, E., Zhang, T., Haugh, K. A., Uchil, P. D., Ochsenbauer, C., Kieffer, C., Kumar, P., Hope, T. J. & Mothes, W. Longitudinal bioluminescent imaging of HIV-1 infection during antiretroviral therapy and treatment interruption in humanized mice. PLoS Pathog 15, e1008161 (2019). PubMed PMC6917343

30 Postler, T. S. & Desrosiers, R. C. The tale of the long tail: the cytoplasmic domain of HIV-1 gp41. J. Virol. 87, 2–15 (2013). PubMed PMC3536369

31 Wyma, D. J., Jiang, J., Shi, J., Zhou, J., Lineberger, J. E., Miller, M. D. & Aiken, C. Coupling of human immunodeficiency virus type 1 fusion to virion maturation: a novel role of the gp41 cytoplasmic tail. J. Virol. 78, 3429–3435 (2004). PubMed PMC371074

32 Murakami, T., Ablan, S., Freed, E. O. & Tanaka, Y. Regulation of human immunodeficiency virus type 1 Env-mediated membrane fusion by viral protease activity. J. Virol. 78, 1026–1031 (2004). PubMed PMC368813

33 Roy, N. H., Chan, J., Lambele, M. & Thali, M. Clustering and mobility of HIV-1 Env at viral assembly sites predict its propensity to induce cell-cell fusion. J. Virol. 87, 7516–7525 (2013). PubMed PMC3700267

34 Krementsov, D. N., Rassam, P., Margeat, E., Roy, N. H., Schneider-Schaulies, J., Milhiet, P. E. & Thali, M. HIV-1 assembly differentially alters dynamics and partitioning of tetraspanins and raft components. Traffic 11, 1401–1414 (2010). PubMed PMC4073295

35 Krementsov, D. N., Weng, J., Lambele, M., Roy, N. H. & Thali, M. Tetraspanins regulate cell-to-cell transmission of HIV-1. Retrovirology 6, 64 (2009). PubMed PMC2714829

36 Nydegger, S., Khurana, S., Krementsov, D. N., Foti, M. & Thali, M. Mapping of tetraspanin-enriched microdomains that can function as gateways for HIV-1. J. Cell Biol. 173, 795–807 (2006). PubMed PMC2063894

37 Khurana, S., Krementsov, D. N., de Parseval, A., Elder, J. H., Foti, M. & Thali, M. Human immunodeficiency virus type 1 and influenza virus exit via different membrane microdomains. J. Virol. 81, 12630–12640 (2007). PubMed PMC2168970

38 Weng, J., Krementsov, D. N., Khurana, S., Roy, N. H. & Thali, M. Formation of syncytia is repressed by tetraspanins in human immunodeficiency virus type 1-producing cells. J. Virol. 83, 7467–7474 (2009). PubMed PMC2708618

39 Symeonides, M., Lambele, M., Roy, N. H. & Thali, M. Evidence showing that tetraspanins inhibit HIV-1-induced cell-cell fusion at a post-hemifusion stage. Viruses 6, 1078–1090 (2014). PubMed PMC3970140

40 Whitaker, E. E., Matheson, N. J., Perlee, S., Munson, P. B., Symeonides, M. & Thali, M. EWI-2 Inhibits Cell-Cell Fusion at the HIV-1 Virological Presynapse. Viruses 11 (2019). PubMed PMC6950393

41 Roy, N. H., Lambele, M., Chan, J., Symeonides, M. & Thali, M. Ezrin is a component of the HIV-1 virological presynapse and contributes to the inhibition of cell-cell fusion. J. Virol. 88, 7645–7658 (2014). PubMed PMC4054451

42 Kumar, R., Barua, S., Tripathi, B. N. & Kumar, N. Role of ROCK signaling in virus replication. Virus Res. 329, 199105 (2023). PubMed PMC10194102

43 Joo, E. & Olson, M. F. Regulation and functions of the RhoA regulatory guanine nucleotide exchange factor GEF-H1. Small GTPases 12, 358–371 (2021). PubMed PMC8583009

44 Zhang, H., Wang, L., Kao, S., Whitehead, I. P., Hart, M. J., Liu, B., Duus, K., Burridge, K., Der, C. J. & Su, L. Functional interaction between the cytoplasmic leucine-zipper domain of HIV-1 gp41 and p115-RhoGEF. Curr. Biol. 9, 1271–1274 (1999). PubMed PMC4513661

45 Douglas, M. E., Davies, T., Joseph, N. & Mishima, M. Aurora B and 14-3-3 coordinately regulate clustering of centralspindlin during cytokinesis. Curr. Biol. 20, 927–933 (2010). PubMed PMC3348768

46 Basant, A., Lekomtsev, S., Tse, Y. C., Zhang, D., Longhini, K. M., Petronczki, M. & Glotzer, M. Aurora B kinase promotes cytokinesis by inducing centralspindlin oligomers that associate with the plasma membrane. Dev. Cell 33, 204–215 (2015). PubMed PMC4431772

47 Ikeda, T., Symeonides, M., Albin, J. S., Li, M., Thali, M. & Harris, R. S. HIV-1 adaptation studies reveal a novel Env-mediated homeostasis mechanism for evading lethal hypermutation by APOBEC3G. PLoS Pathog 14, e1007010 (2018). PubMed PMC5931688

48 Bruce, J. W., Park, E., Magnano, C., Horswill, M., Richards, A., Potts, G., Hebert, A., Islam, N., Coon, J. J., Gitter, A., Sherer, N. & Ahlquist, P. HIV-1 virological synapse formation enhances infection spread by dysregulating Aurora Kinase B. PLoS Pathog 19, e1011492 (2023). PubMed PMC10374047

49 Nakane, S. & Matsuda, Z. Dual Split Protein (DSP) Assay to Monitor Cell-Cell Membrane Fusion. Methods Mol Biol 1313, 229–236 (2015). PubMed 25947669

50 Verhoef, L. G., Mattioli, M., Ricci, F., Li, Y. C. & Wade, M. Multiplex detection of protein-protein interactions using a next generation luciferase reporter. Biochim Biophys Acta 1863, 284–292 (2016). PubMed 26646257

51 Jolly, C. T cell polarization at the virological synapse. Viruses 2, 1261–1278 (2010). PubMed PMC3185707

52 Yuce, O., Piekny, A. & Glotzer, M. An ECT2-centralspindlin complex regulates the localization and function of RhoA. J. Cell Biol. 170, 571–582 (2005). PubMed PMC2171506

53 Opyrchal, M., Allen, C., Msaouel, P., Iankov, I. & Galanis, E. Inhibition of Rho-associated coiled-coil-forming kinase increases efficacy of measles virotherapy. Cancer Gene Ther. 20, 630–637 (2013). PubMed PMC3903406

54 Tieu, K. V., Espey, M., Narayanan, A., Heise, R. L., Alem, F. & Conway, D. E. SARS-CoV-2 S-protein expression drives syncytia formation in endothelial cells. Sci Rep 15, 3549 (2025). PubMed PMC11775288

